# Exploratory re-encoding of Yellow Fever Virus genome: new insights for the design of live-attenuated viruses

**DOI:** 10.1101/256610

**Authors:** R. Klitting, T. Riziki, G. Moureau, G. Piorkowski, E. A. Gould, X. de Lamballerie

## Abstract

Virus attenuation by genome re-encoding is a pioneering approach for generating effective live-attenuated vaccine candidates. Its core principle is to introduce a large number of synonymous substitutions into the viral genome to produce stable attenuation of the targeted virus. Introduction of large numbers of mutations has also been shown to maintain stability of the attenuated phenotype by lowering the risk of reversion and recombination of re-encoded genomes. Identifying mutations with low fitness cost is pivotal as this increases the number that can be introduced and generates more stable and attenuated viruses. Here, we sought to identify mutations with low deleterious impact on the *in vivo* replication and virulence of yellow fever virus (YFV). Following comparative bioinformatic analyses of flaviviral genomes, we categorized synonymous transition mutations according to their impact on CpG/UpA composition and secondary RNA structures. We then designed 17 re-encoded viruses with 100-400 synonymous mutations in the NS2A-to-NS4B coding region of YFV *Asibi* and *Ap7M* (hamster-adapted) genomes. Each virus contained a panel of synonymous mutations designed according to the above categorisation criteria. The replication and fitness characteristics of parent and re-encoded viruses were compared *in vitro* using cell culture competition experiments. *In vivo* laboratory hamster models were also used to compare relative virulence and immunogenicity characteristics. Most of the re-encoded strains showed no decrease in replicative fitness *in vitro*. However, they showed reduced virulence and, in some instances, decreased replicative fitness *in vivo*. Importantly, the most attenuated of the re-encoded strains induced robust, protective immunity in hamsters following challenge with *Ap7M*, a virulent virus. Overall, the introduction of transitions with no or a marginal increase in the number of CpG/UpA dinucleotides had the mildest impact on YFV replication and virulence *in vivo*. Thus, this strategy can be incorporated in procedures for the finely tuned creation of substantially re-encoded viral genomes.

## Introduction

The *Flavivirus* genus (family *Flaviviridae)* includes 53 taxonomically recognized species and at least 19 “tentative” species (1). The flaviviruses display an exceptional diversity of ecological networks that widely correlate with the phylogenetic relationships within the genus (2-5). To date, the majority of recognised flaviviruses are arthropod-borne viruses (arboviruses). They are transmitted amongst vertebrate hosts by mosquitoes (MBFV), ticks or sandflies (TBFV). The genus also includes viruses with no known vector (“NKV”) and some that infect only insects (insect-specific flaviviruses, “ISFV”) (6, 7). Several important (re)-emerging human pathogens fall within the MBFV and TBFV groups, notably dengue virus (DENV), yellow fever virus (YFV), tick-borne encephalitis virus (TBEV), Japanese encephalitis virus (JEV), West Nile virus (WNV) and the recently emerged, Zika virus (ZIKV). Flaviviruses are spherical, enveloped particles (virions) of *ca.* 50nm diameter, incorporating single-stranded positive sense RNA genomes. The capped, 11 kilobase genome includes a single open reading frame (ORF), flanked at its 5’ and 3’ termini by structured, non-coding regions, essential for viral RNA translation and replication (8). ORF translation gives rise to a polyprotein that is cleaved co- and post-translationally into 3 structural (C-prM-E) and 7 non-structural proteins (NS1-2A-2B- 3-4A-4B-5) (9, 10).

YFV is the type species of the genus *Flavivirus*, that owes its name (*flavus* is the Latin word for yellow) to the jaundice associated with the liver dysfunction characteristic of yellow fever disease. In humans, YFV primarily infects the liver, often resulting in severe viscerotropic infections and high fever (11). The severity of YF infections ranges from “inapparent” (*ie.* sub-clinical) to fatal haemorrhagic disease with a mortality rate between 20 and 50% amongst symptomatic cases (12). YFV originated in the tropical African forests where non-human primates (NHP) and a variety of *Aedes* spp. mosquitoes, inhabit the forest canopy, providing the hosts and vectors for viral maintenance through a sylvatic enzootic transmission cycle.

Human-mosquito-human transmission cycles usually arise incidentally when humans encroach on the forest or neighbouring savannah environment and then inadvertently take the virus back to the villages or towns/cities, leading to urban epidemics that can be extensive. The most recent of these large-scale African epidemics happened in Angola and Democratic Republic of Congo in 2015 and 2016 (13, 14).YFV transmission is also maintained in the tropical forests of South-America. However, in this case the sylvatic transmission cycle is epizootic and human cases are primarily “spill-over” events from these sylvatic outbreaks (15-19), as exemplified by the recent epidemic that occurred in Brazil in 2016-2017 (19).

Over the past five or six centuries YFV has killed millions of humans and primates, hence the need for effective methods of prevention. As a response to this need, the live-attenuated vaccine (strain 17D) was developed in 1936 by M.Theiler (20). It was first tested in 1938 in Brazil (21, 22) and two substrains (17DD and 17D-204) are still widely used as the source for vaccine manufacture (23) in Brazil (17DD) and the Old World (17D-204). Throughout decades of use of 17D vaccine, with 20 to 60 million doses distributed annually (24), YF vaccination has proven to be safe and effective, providing long-lasting immunity (25) with rare adverse events (24). The YF 17D strain was obtained through serial passages (>200) of the wild-type (WT) *Asibi* strain in mouse and chicken embryo tissues (25). Similar empirical methods have also been used for the production of notable live-attenuated vaccines against poliovirus (26), measles (27-29) and mumps (30). This strategy relies on attenuation resulting from a relatively limited number of attenuating mutations that are, most often, non-synonymous. However, with all of these live-attenuated viruses there is a risk of (i) attenuation reversion, as described in the case of the poliovirus vaccine (31); (ii) generation of new biological properties, as illustrated by the gain of neurovirulence observed for the YFV French neurotropic vaccine strain (32); and (iii) recombination, as documented for poliovirus (31, 33, 34).

A new, promising, approach for virus attenuation called codon re-encoding, was developed a decade ago by Burns (35) and Mueller (36). Based on the initial observation that usage amongst synonymous codons is highly non-random in the genome of viruses, they hypothesised that this equilibrium could be modified in a manner that should be relatively detrimental to virus replication. They provided the first evidence supporting this concept by introducing slightly detrimental synonymous codons within the encoding region of the poliovirus genome, i.e. without modifying the encoded proteins after which they observed a decrease in viral replication capacity. This codon re-encoding strategy bypasses the limitations of empirical attenuation methods: importantly, the high number of mutations involved decreases the risk of reversion of the attenuating mutations and reduces the likelihood of vaccine strain recombination (with either vaccine or wild-type viruses). The use of silent mutations also lowers the risk of emergence of untoward biological properties.

The first rationale for codon usage bias was that codon abundance correlated with that of isoaccepting tRNAs and could influence the level of protein production within a given host (37). Several other possibilities have been proposed since then, including implication in mRNA structure and its folding (38, 39), microRNA-targeting (40, 41) and enhanced recognition by the immune system (42), that may vary according to the host (43, 44). Specific and random re-encoding approaches have been efficiently applied to several human RNA viruses including poliovirus, influenza A virus, human immunodeficiency virus, respiratory syncytial virus, chikungunya virus (CHIKV), JEV, TBEV and DENV (35, 36, 45-47). Considering a major role of genome-wide mutational processes in the shaping of synonymous sites (48), specific re-encoding approaches include codon and codon-pair deoptimization as well as increase of CpG/UpA dinucleotide frequency. On the other hand, the efficiency of random codon re-encoding strategies for the attenuation of CHIKV *in vitro* and of TBEV *in vivo* suggests an important influence of local constraints (*e.g.* cis-acting sequences, miRNA targeting) on genome encoding (45, 46, 49, 50).

Much remains to be learned to understand the multiple mechanisms that shape synonymous codon usage and its contribution to the attenuation process during re-encoding. In this exploratory work, we defined several types of silent mutations and sought to identify which one(s) had the least detrimental impact on viral replicative fitness. In the future, such silent changes could be introduced in large numbers along the genome of vaccine candidates. YFV provides a suitable experimental model because (i) the virus can be produced conveniently and modified using the Infectious Subgenomic Amplicons (ISA) reverse genetics method (51) and (ii) an excellent animal model of infection exists in juvenile Syrian Golden hamsters, to enable the comparison of wild-type and re-encoded strains *in vivo*. This model was established in our laboratory using the *Asibi*-derived, YF *Ap7M* strain (Klitting *et al.*, accepted manuscript)derived from the previously described hamster-virulent YF *Asibi*/hamster p7 strain (47). When inoculated into hamsters, both strains induce a disease with features close to that of the human disease.

Starting with a bioinformatic analysis of flaviviral genomes, we defined several types of mutations and used them for the design of 8 re-encoded sequences. The re-encoded viruses were derived from the wild-type *Asibi* strain and the hamster-adapted *Ap7M* strain, with which they were compared both *in vitro* and *in vivo* through fitness assays, infection assays and complete genome deep-sequencing. Although only the most heavily re-encoded virus (741 mutations) showed an observable decrease in replicative fitness *in vitro*, a range of attenuation levels was observed for the re-encoded variants when tested *in vivo*. The lowest impact was observed for strains that were re-encoded with transitions that did not include new CpG/UpA dinucleotides. In addition, we showed that the strains with the most attenuated phenotypes *in vivo* could induce robust protective immunity in hamsters following challenge with the lethal YF *Ap7M* virus.

## Results

*The results of the in silico analysis that served as a basis for the design of the re-encoded strains are provided in the Supplementary Results S2 section.*

### Design strategies for re-encoded strains

#### Framework design

Previous analyses on the impact of random codon re-encoding on chikungunya (CHIKV) and tick-borne encephalitis (TBEV) viruses, were used as a starting point to obtain observable *in vitro* and/or *in vivo* attenuation. Nougairède and colleagues observed a decrease in CHIKV replicative fitness *in vitro*, using 264 to 882 mutations located within the NSp1, NSp4 and/or envelope protein encoding regions (45). Clear *in vivo* attenuation of TBEV was achieved by de Fabritus and colleagues using as few as 273 mutations located in the NS5 coding region (46).

Based on those results, we defined a framework for the design of YFV re-encoded strains. For each variant, we used a maximum of 350 synonymous mutations (*ca.* 1 change for every 10 nucleotides). We chose to re-encode the region located within the coding sequences of proteins NS2A to NS4B, so that re-encoding would not affect the structural proteins, the viral RNA-dependent RNA polymerase, or the highly structured, 5’ and 3’ ends of the genome. The highly re-encoded variant rTs4 was reconstructed by the addition of 388 synonymous mutations in the NS5 encoding region. In all cases, the structural proteins encoding sequence remained unchanged, which was an advantage as the production of hamster-virulent YFV variants required the use of a modified E protein sequence. To lower the risk of major detrimental effects of re-encoding (*e.g.* disruption of cis-acting sequences), we analysed an alignment of 35 YFV complete coding sequences (CDS) and defined only synonymous sites for which at least one mutation could be observed as eligible for re-encoding. These will be referred to as “editable sites”. Finally, we avoided creating or removing any rare codons during the re-encoding process (see definition in Supplementary Protocols S1).

#### Re-encoding strategies

Following *in silico* analysis of flaviviral genomes, we defined two major types of synonymous mutations (see below). The creation of additional TCG trinucleotide patterns was considered a “specific” type of mutation. In the genus *Flavivirus*, TCG trinucleotides are frequent in Insect-Specific Flavivirus (ISF) genomes and sparse in both No Known Vector virus (NKV) and arbovirus genomes (see Supplementary results). In addition, this pattern may enhance the innate immunity in vertebrate cells, as suggested by Greenbaum, Jimenez-Baranda and colleagues (43, 53). “Specific” mutations also included the creation of CpG and UpA dinucleotides, which were the least frequently introduced dinucleotides (CpG and UpA) in YFV species (see Supplementary results). This is in accordance with the previous observation of dinucleotide bias in the genomes of many viruses, in a fashion that was suggested to, at least partly, mirror the dinucleotide usage of their host(s) (54, 55). In vertebrates, CpG dinucleotide bias is commonly seen as the consequence of cytosine methylation-deamination (56-58) and the detection of such patterns may be a part of the host-defence mechanism against non-self RNA (59-61). The introduction of CpG and UpA dinucleotides has been frequently used in re-encoding studies (42, 62-64). By contrast, “non-specific” types of mutations did not cause any increase in CpG/UpA dinucleotides. This category includes transitions located on and outside sites associated with the putative secondary structures identified in the YFV genome during *in silico* analysis (see Supplementary results). All mutation types defined in this study are described in Table 1.

**Table 1.** Mutation categories

|  | Non-specific | Specific |
| --- | --- | --- |
| Mutations | -Simple transitions<br>-Transitions on sites associated with putative secondary structures | - CpG/UpA dinucleotide transitions<br>- TCG trinucleotide introductions |

Note: Based on bioinformatic analysis, 2 major types of mutation were defined.

#### Design

In total, 9 re-encoded viruses were designed based on the *Asibi* strain sequence. Aside from the highly re-encoded rTs4 virus, all Asibi-derived re-encoded viruses were designed with 100 to 400 mutations located between positions 3,924 and 6,759 of YF *Asibi* complete coding sequence (CDS). First, YFV rTs3 was designed by introducing exclusively “simple” transitions (siTs) into the target region, within the CDS of the reference strain *Asibi* (AY640589). The term “simple” refers to synonymous, non-specific (no CpG/UpA introduction) mutations located outside putative secondary structures. YFV rTs3 included 353 substitutions, i.e., the highest possible number that could be introduced into the target region. All other re-encoded viruses except YFV rTCG were designed starting from the rTs3 virus by (i) reverting some of the siTs back to the original sequence (rTs1, rTs2), or (ii) substituting some of the siTs for a different type of mutation (rUA, rCG, rSS and rN). The rationale for this design procedure was to obtain viruses that could be compared one to another, as they would share a common background of mutations.

Overall the 9 initial re-encoded viruses were designed as follows:

- YFV rTs1, rTs2 and rTs3, include increasing numbers of “simple” transitions (siTs).
- YFV rSS includes non-specific transitions on sites corresponding to putative secondary structures (SII-Ts).
- YFV rUA and rCG include transitions leading to an increase in the number of UpA or CpG dinucleotides into the viral sequence (UA- and CG-Ts), respectively.
- YFV rN includes both transitions and transversions, some of which lead to the introduction of 83 UpA and 54 CpG dinucleotides into the viral sequence.
- YFV rTCG includes transitions leading to an increase in the number of TCG dinucleotides into the viral sequence (TCG-Ts). Importantly, this strain was not derived from rTs3 and exclusively includes TCG-Ts.
- YFV rTs4 includes 388 additional siTs that were introduced into the genome of rTs3 virus, between positions 6,846 and 9,765 (in CDS).

Finally, 8 hamster-adapted re-encoded strains were created from the *Asibi*-derived variants (other than the heavily re-encoded rTs4 virus) by introducing exactly the same mutations in the CDS of the hamster-virulent, YFV *Ap7M* virus. For all re-encoded viruses, the design is further detailed in the “Material and Methods” section.

Following re-encoding, the GC% and Effective Number of Codons (ENC) were calculated for each sequence and compared to those obtained from a dataset of 35 YFV genome sequences (non-vaccine strains) using SSE software (v1.3) (65). Apart from the most re-encoded, rTs4 (GC%:52,3%; ENC:52,1), all the other strains fell within the YFV species range in terms of both GC% (48,9%-50,5%) and ENC (52,4 - 54,7). Details regarding re-encoded strain design are available in Table 2.

**Table 2.** Asibi and Ap7M-derived re-encoded strain description

| Name | Description | Transitions | Transversions | CpG-Ts | UpA-Ts | SII-Ts | TCG-Ts | NE sites-Ts | Total | GC content (%) | ENC | Titre (TCID50/mL) | Name (*Ap7M*-derived viruses) | Titre (TCID50/mL) |
| --- | --- | --- | --- | --- | --- | --- | --- | --- | --- | --- | --- | --- | --- | --- |
| ***Asibi*** | *Asibi* | 0 | 0 | 0 | 0 | 0 | 0 | 0 | 0 | 49,7 | 52,8 | 1E+08 | ***Ap7M*** | 9E+08 |
| ***rTs1*** | siTs-low | 101 | 0 | 0 | 0 | 0 | 2 | 0 | 101 | 49,7 | 52,9 | 4E+05 | ***rTs1h*** | 6E+08 |
| ***rTs2*** | siTs-intermediate | 247 | 0 | 0 | 0 | 0 | 2 | 0 | 247 | 49,5 | 52,7 | 4E+07 | ***rTs2h*** | 6E+08 |
| ***rTs3*** | siTs-high | 353 | 0 | 0 | 0 | 0 | 3 | 0 | 353 | 49,4 | 52,6 | 4E+06 | ***rTs3h*** | 6E+08 |
| ***rUA*** | UpA-Ts | 326 | 0 | 0 | 59 | 0 | 2 | 0 | 326 | 49,6 | 53,5 | 3E+06 | ***rUAh*** | 4E+08 |
| ***rCG*** | CpG-Ts | 325 | 0 | 59 | 0 | 0 | 19 | 0 | 325 | 49,7 | 52,8 | 3E+06 | ***rCGh*** | 4E+08 |
| ***rSS*** | Ts on putative secondary structures | 339 | 0 | 0 | 0 | 50 | 3 | 0 | 339 | 49,7 | 52,9 | 7E+06 | ***rSSh*** | 4E+08 |
| ***rN*** | Combinatory strategy | 179 | 54 | 59 | 83 | 0 | 17 | 0 | 233 | 49,5 | 52,7 | 4E+06 | ***rNh*** | 6E+06 |
| ***rTCG*** | Ts to TCG | 46 | 72 | 111 | 0 | 12 | 118 | 11 | 118 | 49,4 | 52,6 | 2E+06 | ***rTCGh*** | 6E+07 |
| ***rTs4*** | siTs-very high | 741 | 0 | 0 | 0 | 0 | 7 | 0 | 741 | 52,3 | 52,1 | 6E+03 | - | - |

Note: For each YFV re-encoded strain, the number of mutations introduced within the CDS of the reference strain *Asibi* (Genbank AN: AY640589) are given both as a total and with details according to the type of mutation used. The following abbreviations were used: CpG-Ts: transition leading to CpG dinucleotide introduction, UpA-Ts: transition leading to UpA dinucleotide introduction; SII-Ts: transition on sites corresponding to putative secondary structures; NE sites-Ts: transition on sites for which no mutation has been reported in the 35 YFV CDS alignment. G+C content was calculated on viral CDS and is presented as percentage. For each viral CDS, the ENC was calculated using SEE software (v1.3) (64). Infectious titres were determined on BHK21 cells and are presented as TCID50/mL for both *Asibi*-derived viruses and the corresponding, *Ap7M*-derived viruses.

### *In vitro* behaviour of YFV re-encoded strains

*All the strains used in this study (1 WT, 1 hamster-adapted and 17 re-encoded) were produced using the ISA method, as detailed in Fig 1 and in the “Material and Methods” section.*

**Figure 1.**
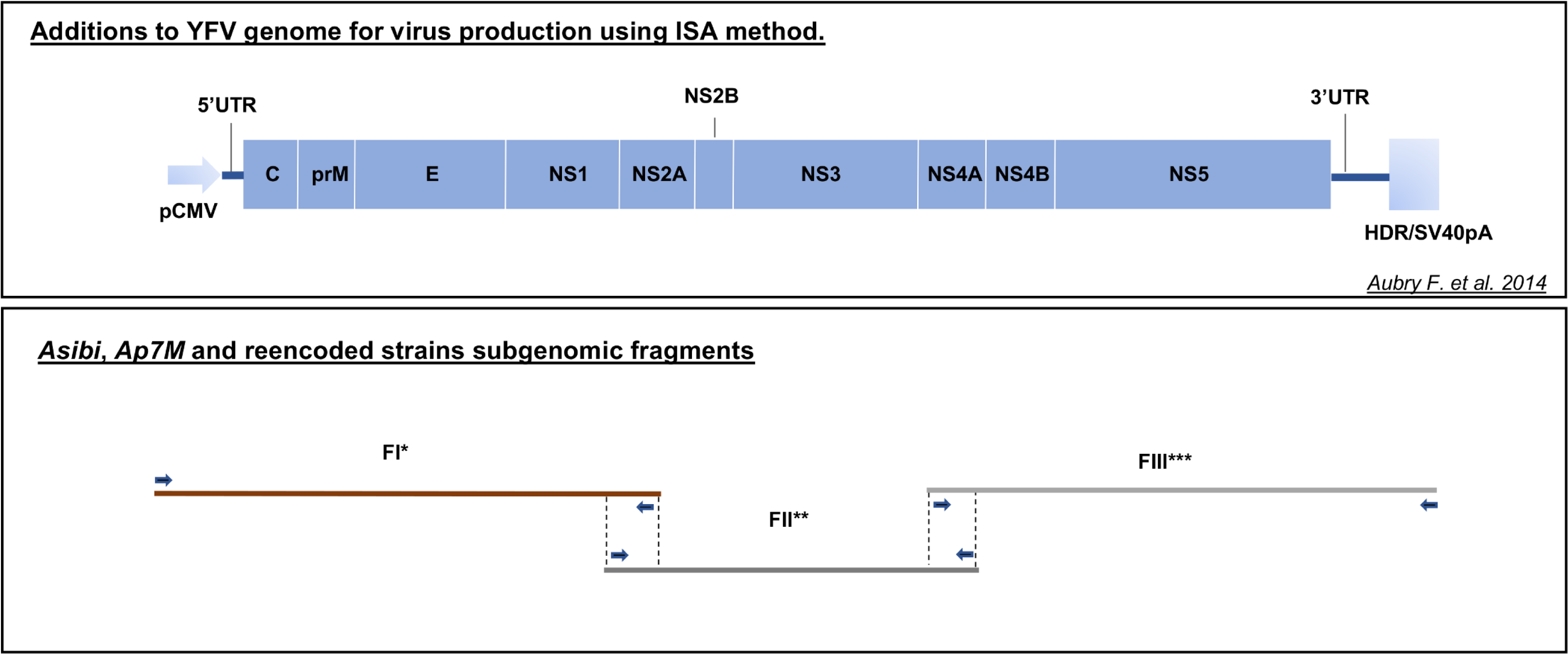
*Subgenomic fragments used for wild-type and re-encoded virus production using the Infectious Subgenomic Amplicon method.* *For *Ap7M* strain and hamster-adapted re-encoded strain (*Ap7M*-derived) production, the subgenomic fragment FIAp7 was amplified from the corresponding plasmid and combined to subgenomic fragments FII (wild-type or re-encoded) and FIII (wild-type). **For each of the YFV re-encoded strains (both *Asibi* and *Ap7M*-derived) production, a different subgenomic fragment FII was amplified from the corresponding plasmid. ***For the production of YFV re-encoded strain rTs4, the subgenomic fragment FIIIR was amplified from the corresponding plasmid and combined to fragments FI (wild-type) and FII (re-encoded).

#### Sequence analysis

Next Generation Sequencing (NGS) was performed on the cell culture supernatant media (viral stocks *i.e.*, after 3 passages). For all viruses, both the integrity of the genome and the intra-population genetic diversity were assessed on the CDS. At the level of intra-population genetic diversity, a variant subpopulation (further referred to as “variant”) was regarded as major when its corresponding nucleotide proportion (CNP) was over 75%. Only consensus mutations associated with major variants were considered for further analysis. All sequencing results are shown in Tables 3 and 4.

**Table 3.** Sequence analysis of Asibi and Asibi-derived re-encoded viruses

| Strain name | Re-encoded region | Position (nt) | Protein | Consensus mutation | Expected sequence (nt) | Observed sequence (1<sup>st</sup> variant (nt)) | Observed sequence (2<sup>nd</sup> variant (nt)) | 1<sup>st</sup> variant proportion | 2<sup>nd</sup> variant proportion |
| --- | --- | --- | --- | --- | --- | --- | --- | --- | --- |
| *Asibi* | - | - | - | - | - | - | - | - | - |
| *rTs1* | 3927-6753 | - | - | - | - | - | - | - | - |
| *rTs2* | 3927-6753 | 5430 | - | - | C | T | C | 77,5% | 22,3% |
|  |  | 5442 | - | - | C | T | C | 82,0% | 18,0% |
| *rTs3* | 3927-6753 | 4357 | NS2B | M99V | A | G | A | 78,3% | 21,7% |
|  |  | 9150 | - | - | A | G | A | 78,5% | 21,5% |
| *rUA* | 3927-6753 | - | - | - | - | - | - | - | - |
| *rCG* | 3924-6750 | 4396 | NS2B | F112L | T | C | T | 88,2% | 11,8% |
|  |  | 5430 | - | - | C | T | - | >95% | - |
|  |  | 5667 | - | - | A | G | - | >95% | - |
|  |  | 6082 | NS3 | C544R | T | C | - | >95% | - |
| *rSS* | 3939-6759 | 561 | - | - | T | C | - | >95% | - |
|  |  | 3190 | NS1 | I286V | A | G | - | >95% | - |
|  |  | 4308 | - | - | G | T | - | >95% | - |
|  |  | 4623 | - | - | A | G | - | >95% | - |
|  |  | 6012 | - | - | A | G | - | >95% | - |
| *rN* | 3924-6750 | - | - | - | - | - | - | - | - |
| *rTCG* | 3939-6759 | 2468 | NS1 | I44T | T | C | - | >95% | - |
|  |  | 2795 | NS1 | Q154R | A | G | - | >95% | - |
|  |  | 7629 | - | - | T | C | - | >95% | - |
|  |  | 8565 | - | - | T | C | - | >95% | - |
|  |  | 8916 | - | - | T | C | - | >95% | - |
|  |  | 9297 | - | - | T | C | - | >95% | - |
| *rTs4* | 3927-9765 | - | - | - | - | - | - | - | - |

Note: For each virus, the consensus changes found in the CDS from the viral stock are detailed above. The expected sequence refers to that of plasmids that served for subgenomic fragment amplification while producing viruses using the ISA method. Only consensus mutations associated with major variants (CNP>75%) are described. Amino Acid (AA) positions are given with reference to the beginning of mature protein sequence (ie. AA position 107 in prM, written prM 107, corresponds to AA position 228 within the precursor polyprotein). A detailed map of protein positions within the YFV polyprotein sequence is provided in the supplementary Table S11.

**Table 4.** Sequence analysis of Asibi, and both Asibi and Ap7M-derived re-encoded viruses

| **VIRUS** | **Consensus mutations** | **Consensus NS mutations** | **Variants** | **Reversions** | **Consensus mutations in re-encoded region** | **Variants in re-encoded region** | **Total no. reads** | **Coverage depth (x)** | **Mean coverage/site** |
| --- | --- | --- | --- | --- | --- | --- | --- | --- | --- |
| *Asibi* | 0 | 0 | 12 | 0 | 0 | 1 | 317220 | 30 | 6006 |
| *rTs1* | 0 | 0 | 12 | 0 | 0 | 0 | 286332 | 27 | 5521 |
| *rTs2* | 2 | 0 | 9 | 2 | 2 | 3 | 410433 | 40 | 7625 |
| *rTs3* | 2 | 1 | 6 | 0 | 1 | 2 | 167768 | 16 | 3208 |
| *rUA* | 0 | 0 | 16 | 0 | 0 | 5 | 113558 | 11 | 2075 |
| *rCG* | 4 | 2 | 5 | 2 | 4 | 1 | 419297 | 40 | 7900 |
| *rSS* | 5 | 1 | 1 | 0 | 3 | 0 | 55624 | 5 | 1026 |
| *rN* | 0 | 0 | 11 | 0 | 0 | 2 | 223184 | 21 | 4274 |
| *rTCG* | 6 | 2 | 4 | 0 | 0 | 1 | 249701 | 24 | 4580 |
| *rTs4* | 0 | 0 | 15 | 0 | 0 | 9 | 176219 | 17 | 3227 |
| *rTs1h* | 0 | 0 | 13 | 0 | 0 | 4 | 619634 | 60 | 10949 |
| *rTs2h* | 0 | 0 | 22 | 0 | 0 | 6 | 144211 | 14 | 2442 |
| *rTs3h* | 3 | 3 | 13 | 0 | 1 | 3 | 528403 | 51 | 11304 |
| *rUAh* | 0 | 0 | 17 | 0 | 0 | 5 | 159007 | 15 | 2724 |
| *rCGh* | 0 | 0 | 10 | 0 | 0 | 0 | 297217 | 29 | 6220 |
| *rSSH* | 0 | 0 | 19 | 0 | 0 | 2 | 256125 | 25 | 4258 |
| *rNh* | 5 | 1 | 4 | 0 | 0 | 0 | 514717 | 50 | 10693 |
| *rTCGh* | 2 | 2 | 7 | 0 | 0 | 0 | 163170 | 15 | 2413 |

Note: Counts of consensus mutations and variant subpopulations are presented as raw numbers. Consensus reversion mutations were defined with reference to YFV strain *Asibi*. The coverage depth has been calculated given that the total length of YFV CDS is 10,236 nt. No variant was taken into account with a coverage lower than 1000 reads. No consensus mutation was taken into account with a coverage lower than 50 reads.

During virus production, sequence variability could arise as an artefact of the production method itself (PCR amplification of DNA fragments) and as the result of the adaptation of re-encoded strains to culture conditions.

No consensus mutation associated with a major variant was identified in the genome of the parent strain YFV *Asibi*, which only included twelve minor variants. In re-encoded genome sequences, low numbers of consensus mutations were observed, ranging from 0 to 6, with a mean of 2 consensus changes/virus and 39% of non-synonymous (NS) mutations on average. Consensus mutations were equally distributed between re-encoded and non-re-encoded regions and, nearly no convergence was observed between viral sequences. Only two viral sequences (YFV rTs2 and rCG) had consensus mutation reversions. One was common to both strains and corresponded to a siTs mutation (C to T), at CDS position 5,430. The two others were also siTs mutation reversions (C to T and A to G, respectively), located at CDS positions 5,442 and 5,667 of YFV rTs2 and rCG, respectively. The absence of consensus NS change in the sequence of the original *Asibi* strain and the very limited convergence between viral sequences are in accordance with the low number (3) of *in cellulo* passages. This strongly suggests that the consensus mutations observed in viruses YFV rTs3, rCG, rSS and rTCG did not result from selective pressure due to culture conditions but may be artefacts of the production method.

The total numbers of variants, per virus, ranged from 1 to 16, with a mean of 9 variants per virus. Interestingly, variants were most frequently observed outside the re-encoded regions (75% of variants outside the re-encoded regions), similar to what was observed for the parental sequence, (92% of variants outside the re-encoded region). The low number of reversions and the limited sequence variability at re-encoded sites accord with previous work showing that re-encoded CHIKV strains evolved mostly through compensatory mutations rather than mutation reversions (45).

#### *In vitro* replicative fitness

The infectious titre was estimated using a TCID_50_ assay in BHK21 cells for both *Asibi* and *Asibi*-derived re-encoded strains using infectious cell supernatant medium (viral stock) (results detailed in Table 2). The maximal infectious titre was observed for the wild-type strain *Asibi* (10^8^ TCID_50_/mL). A 1 to 2-log decrease was observed for most re-encoded strains, with infectious titres ranging between 2.10^6^ and 4.10^7^ TCID_50_/mL. However, the least re-encoded virus, YFV rTs1 (101 mutations), was unexpectedly endowed with a remarkably low infectious titre (4.10^5^ TCID_50_/mL) whilst the most re-encoded virus, YFV rTs4 (741 mutations), exhibited the lowest infectious titre (6.10^3^ TCID_50_/mL).

For each of the re-encoded viruses, the replicative fitness was compared to that of the wild-type strain using competition assays. Specific virus detection was achieved for one in two passages using next generation sequencing (NGS) methods by amplifying a 256 nucleotide region and counting virus-specific patterns within the read population. For all re-encoded viruses except rTs4 (≤ 353 mutations), replicative fitness *in vitro* was comparable to that of the *Asibi* strain: in all competitions, both competing viruses could still be detected at the 10^th^ passage (see Fig 2). By contrast, the highly re-encoded rTs4 virus (741 mutations) showed a clear reduction in replicative fitness and could not be detected after the 5^th^ passage.

**Figure 2.**
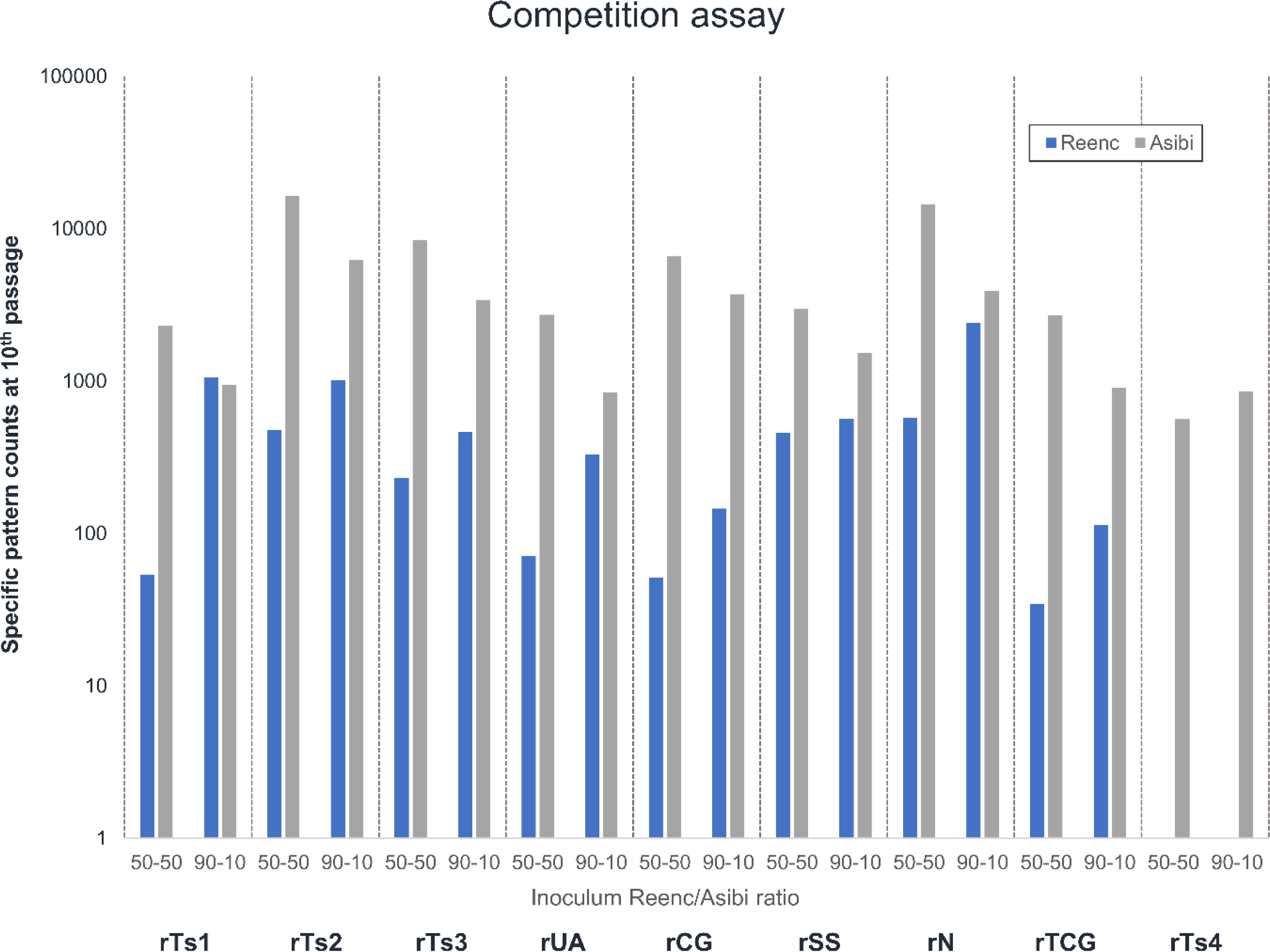
*Competition assays: Wild-type and re-encoded-specific patterns counts at 10th competition passage.* For each WT/re-encoded virus ratio and each re-encoded virus, pattern counts were calculated as a mean for the 3 replicates. For each replicate, the pattern count is the mean count for the three virus-specific patterns. Each specific pattern count is the sum of the sense and reverse patterns counts within a given read population (obtained through NGS sequencing).

### *In vivo* behaviour of YFV re-encoded strains

The infectious titres of viral stocks for *Ap7M* and *Ap7M*-derived re-encoded strains were established in BHK21 cells, using a TCID_50_ assay (results detailed in Table 2). For the majority of re-encoded viruses, no notable difference was observed with the parent strain *Ap7M* (9.10^8^ TCID50/mL), with infectious titres ranging from 4.10^8^ to 6.10^8^ TCID50/mL. However, viruses YFV rNh and rTCGh showed a 1 to 2-log reduction in viral titre (6.10^6^ and 6.10^7^ TCID50/mL, respectively).

#### Sequence analysis

NGS was performed on the viral stocks for all *Ap7M*-derived re-encoded viruses as described in the previous section (results shown in Tables 4 and 5).

**Table 5.** Sequence and intra-population diversity in viral culture supernatant media and hamster liver samples for Ap7M and Ap7M-derived re-encoded strains

| Strain name | Position (nt) | Protein | Consensus mutation | Expected sequence (nt) | Observed sequence (consensus (nt)) | AA change | Observed sequence (Inoculum (nt)) |  | Observed sequence (Hamster (nt)) |  |  |
| --- | --- | --- | --- | --- | --- | --- | --- | --- | --- | --- | --- |
|  |  |  |  |  |  |  | 1<sup>st</sup> variant | 2<sup>nd</sup> variant | Retrieved sequence | 1<sup>st</sup> variant | 2<sup>nd</sup> variant |
| *rTs1h* | 836 | prM | A158V | C | T | A → V | C | - | 1-4806, 5430-7566 | T | - |
|  | 4574 | NS3 | G41A | G | C | G → A | G | - |  | C | g |
|  | 5442 | - | - | C | T | - | C | - |  | T | - |
| *rTs2h* | - | - | - | - | - | - | - | - | 1-3858, 4653-5771, 6083-10236 | - | - |
| *rTs3h* | 1649 | Env | T265I | C | T | T → I | C | t | 1-10236 | T | - |
|  | 4604 | NS3 | R51M | T | G | R → M | G | t |  | T | g |
| *rUAh* | - | - | - | - | - | - | - | - | 1-6210, 6492-7565 | - | - |
| *rCGh* | 283 | - | - | T | C | - | T | - | 1-4652, 5433-7564 | C | t |
|  | 1051 | Env | T66A | A | G | T → A | A | - |  | G | a |
|  | 2541 | - | - | A | G | - | A | - |  | G | - |
| *rSSh* | 5439 | - | - | G | A | - | G | - | 14-10228 | A | - |
| *rNh* | 818 | prM | I152T | T | C | I → T | C | - | 7-5236, 5422-10229 | C | - |
|  | 1686 | - | - | T | C | - | C | - |  | C | - |
|  | 2052 | - | - | G | A | - | A | - |  | A | - |
|  | 7490 | NS4B | N241S | A | G | N → S | g | a |  | G | - |
|  | 8253 | - | - | G | A | - | A | g |  | A | - |
|  | 9783 | - | - | A | G | - | G | a |  | G | - |
| *rTCGh* | 2222 | Env | I456T | T | C | I → T | C | t | 11-10223 | C | t |
|  | 7107 | NS4B | I113M | A | G | I → M | G | a |  | G | a |

Note: For each virus, the consensus changes found either in the complete coding sequence from the viral stock or in partial sequences from infected hamster liver homogenates are detailed above. The expected sequence refers to that of plasmids that served for subgenomic fragment amplification while producing viruses using the ISA method. For 1<sup>st</sup> and 2<sup>nd</sup> variant descriptions, capital letters correspond to variant proportions above 75%, lowercase letters to variant proportions between 6 and 75% and dashes to variant proportions below 5%. Amino Acid (AA) positions are given with reference to the beginning of mature protein sequence (i.e. AA position 107 in prM, written prM 107, corresponds to AA position 228 within the precursor polyprotein). A detailed map of protein positions within the YFV polyprotein sequence is provided in the supplementary Table S11.

As observed for *Asibi*-re-encoded viruses, the number of consensus mutations associated with major variants were low, ranging from 0 to 5, with a mean number of 1 consensus change/virus and 59% of non-synonymous mutations on average. Consensus mutations were mainly distributed outside re-encoded regions (90%), with no reversion. As observed for *Asibi*, the parent *Ap7M* strain showed no NS consensus sequence changes associated with major variants. Moreover, the re-encoded viruses which were given a limited number of *in cellulo* passages did not exhibit any evidence of convergent evolution. Based on this evidence, the consensus mutations observed in YFV rTs3h, rNh and rTCGh are likely to be artefacts of the production method rather than the result of adaptation of re-encoded viruses to culture conditions. Notably, for YFV rTs3h, the consensus NS change observed in the inoculum was not essential for virus survival *in vivo* as it was not maintained following infection in hamsters.

The total number of variants per virus varied among strains, ranging from 4 to 22, with a mean number of 13 variants per virus. Similar to what was observed for *Asibi*-derived re-encoded strains, the diversity at re-encoded sites was low, with only a minority of variants being detected within the re-encoded regions (19%).

#### *In vivo* phenotype: comparative study of pathogenicity

Groups of 12 three-week-old female hamsters were inoculated intra-peritoneally with 5.10^5^ TCID_50_ of virus (either *Ap7M* or *Ap7M*-re-encoded strains) and a control group of 2 uninfected hamsters was maintained for weight monitoring. Clinical follow-up included (i) clinical manifestations of the disease (brittle fur, dehydration, prostration and lethargy), (ii) body weight (weight evolution was expressed as a normalised percentage of the initial weight (%IW)) and (iii) death. Three and four hamsters were euthanised at days 3 and 6 post-infection (dpi), respectively, to conduct virology investigations from liver samples, whilst the other five in each group were retained for evaluating mortality rate (endpoint in case of survival: 16 dpi). Due to an issue in group dispatching, in the case of rTs2h strain, 3 hamsters were euthanised at 6 dpi while 6 hamsters were used for mortality rate evaluation. Virology follow-up was achieved by performing qRT-PCR and next generation sequencing on RNA extracted from the liver homogenates. RNA samples were subsequently pooled to obtain one sequence for each virus, reflecting genetic diversity amongst the 5 hamsters kept until death/euthanasia. For all viruses, the *in vivo* phenotype and the sequencing results are detailed in Fig 3 and Tables 4 and 5.

**Figure 3.**
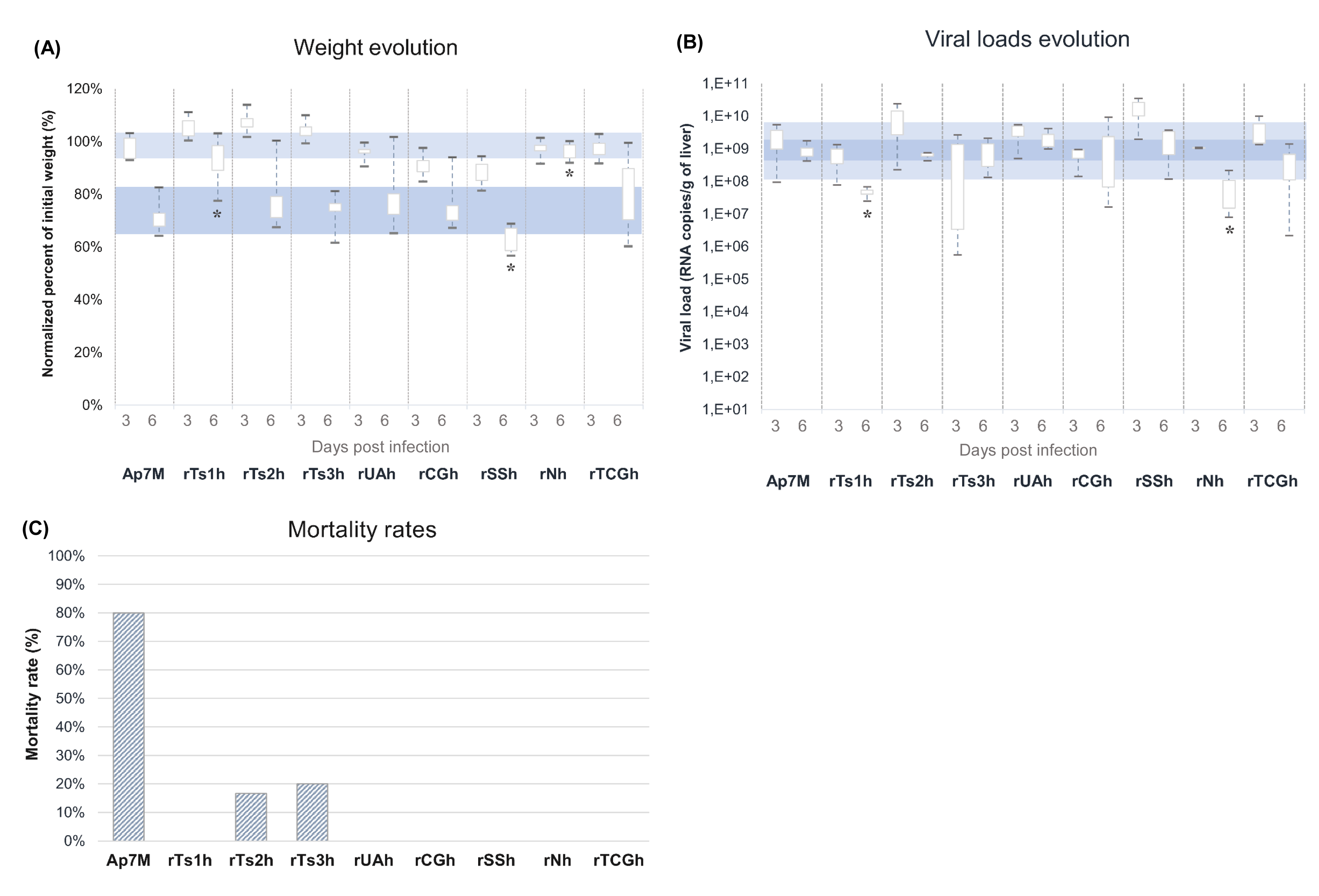
*In vivo phenotypes of Ap7M and Ap7M-derived re-encoded viruses*. Normalized percentages of initial weight (%IW) (A), viral loads at 3/6 dpi (B), as well as mortality rates (C), are illustrated for all viruses. %IW were evaluated on groups of 12 and 9 hamsters at 3 and 6 dpi, respectively. Viral loads were evaluated on groups of 3 and 4 hamsters at 3 and 6 dpi, respectively. Of note, for YFV rTs2h, only 3 hamsters were used for viral load determination at 6 dpi. Mortality rates were evaluated on groups of 5 hamsters for all viruses, apart from rTs2h, for which 6 hamsters were used. Computations of %IW and viral loads are detailed in the corresponding paragraphs within the Materials and Methods section. Significant %IW and viral load differences observed between *Ap7M* and the other viruses at 6 dpi (Wilcoxon rank-sum test p-value <0.05) are indicated by a star (*). The ranges corresponding to viral load values recorded at 3 and 6 dpi during *Ap7M* infection are highlighted by light (3 dpi) and dark (6 dpi) blue rectangles.

Nearly all hamsters inoculated with *Ap7M* strain developed outward signs of illness such as ruffled fur, prostration, dehydration and lethargy. One hamster did not show any sign of illness, both its liver and blood were tested negative for YFV genomes and it was excluded from analysis. A high mortality rate (80%) was observed, similar to previous observations (52). Clinical signs of the disease appeared as early as 4 dpi and all animals died within 2/3 days after onset of the symptoms. Weight loss was not observed at 3 dpi (mean of %IW: 97%), but between 3 and 6 dpi, significant weight losses were recorded (mean of %IW: 71%, Wilcoxon rank sum test, p-value = 0,00032). All livers were found to be YFV-positive by qRT-PCR, with viral RNA loads ranging between 6.10^7^ and 6.10^9^ RNA copies per gram of liver and no significant difference between the viral yields at 3 and 6 dpi (Wilcoxon rank sum test, p-values >0,05). No consensus change was detected in the viral sequence after propagation in hamsters.

A reduction in mortality rate was observed in all groups infected with re-encoded viruses. For most (YFV rTs1h, rUAh, rCGh, rSSh, rNh and rTCGh), the reduction was significant (Logrank test, p-values=0,03960 in all cases), as no hamsters died. However, for YFV rTs2h and rTs3h, mortality was reduced but still observed (17 and 20%, respectively; Logrank test, p-values=0,05820 and 0,07186).

Regardless of the observed mortality rate, clinical signs of illness were observed with varying degrees of severity amongst groups. No association was observed between mortality and weight evolution at 3 or 6 dpi, with non-lethal viruses inducing limited (rNh, rTs1h) as well as severe weight loss (rCGh, rSSh) in infected hamsters. At 3 dpi, weight evolution was heterogeneous but not significantly different from what was observed in *Ap7M*-infected group (Wilcoxon rank sum test, p-values>0,05). An increase up to 7% was observed in YFV rTs1h, rTs2h and rTs3h-infected groups, a slight decrease (between 3 and 4%) in groups infected with YFV rUAh, rNh and rTCGh, and a greater decrease up to 11%, in YFV rCGh and rSSh-infected groups. At 6 dpi, an important weight loss was observed in most groups (up to 38%) except rTs1h and rNh-infected groups, in which significantly higher %IW values were recorded (8 and 4%, respectively, (Wilcoxon rank sum test, p-values=0.00088 and 0,00062, respectively)).

An increase of viral yields at 3 and 6 dpi was not associated with mortality rates. At 3 dpi, viral loads in the liver were close to values observed with YF *Ap7M*, with values ranging from 7.10^8^ to 1.10^10^ RNA copies/gram of liver. At 6 dpi, viral loads remained close to those observed with *Ap7M*. However, in some cases (YFV rTs1h and rNh), significantly lower viral loads were observed (means of 5 and 8.10^7^, respectively. Wilcoxon rank sum test, p-value= 0,03038 in both cases).

Several possibilities can explain the apparent absence of association between mortality and the evolution of viral loads. First, significant reduction in viral loads were always associated with a complete loss of lethality (for viruses rNh and rTs1h), hence they were, to a reasonable extent, associated to pathogenesis. Furthermore, in terms of replicative fitness and thus, of viral loads, the threshold that corresponds with the loss of lethality may be subtle and undetectable under the experimental conditions used here. In addition, the viral loads that are reported here are RNA copy numbers and may not reflect the actual amount of infectious particles. Furthermore, they have been evaluated from liver samples and may be less sharply correlated with disease evolution than viral loads in the blood. Also, variation between individual animals will contribute to the final outcome of the data. Finally, during YFV infection, it is likely that the pathogenesis involves an immunological component (21, 66, 67). Hence, levels of pro- and anti-inflammatory cytokines may be more closely associated with mortality than viral loads and weight evolution. Such criteria were not included in the experimental design.

#### Evaluation of the phenotypic impact of re-encoding

Some conclusions can be drawn by analysing the *in vivo* phenotypes of re-encoded viruses in light of the sequencing data obtained from infected hamsters liver samples (see details in Table 5). For example, the sequences obtained from the livers of hamsters infected with the control strain *Ap7M* did not include consensus mutations. This indicates that the sequence of the parent virus was stable under the experimental conditions and that adaptative mutations were not required for efficient replication *in vivo*.

#### Impact of “non-specific” re-encoding strategies

In general, low detrimental effects resulted from the introduction of simple transitions (siTs) (*i.e.*, with no modification of predicted secondary structures) on the *in vivo* phenotype of YFV. Strain rTs2h (247 siTs) was associated with mild attenuation *in vivo,* with a decreased mortality rate and a reduction in hamster weight loss at 6 dpi but no significant change in the production of viral RNA (Wilcoxon rank sum, p-values>0,05). As the viral sequence remained unchanged during the course of the experiments, the *in vivo* phenotype can be interpreted as being the result of genome re-encoding. rTs3h (353 siTs) and rTs1h (101 siTs) viruses, respectively showed mild and strong attenuation *in vivo*. For both, consensus NS mutations associated with major variants arose during the *in vivo* infections possibly resulting from adaptation of the re-encoded viruses to the *in vivo* conditions during the infection. However, they did not correspond with the recovery of an *in vivo* phenotype similar to that of the parent strain *Ap7M*. Surprisingly, the least re-encoded strain, rTs1h (101 siTs) was more attenuated *in vivo* than the rTs2h strain, that included all rTs1h mutations plus 146 additional siTs. We observed a mild detrimental impact following the introduction of transitions at sites involved in predicted RNA secondary structures. The sequence of strain rSSh involved siTs as well as transitions located on predicted secondary RNA structures and remained stable throughout the *in vivo* experiments. The variant showed a partially attenuated phenotype *in vivo* (complete loss of mortality, no alleviation in weight loss or viral loads) that can thus be attributed to this second re-encoding strategy. Overall, these results illustrate the relative efficiency of the use of “non-specific” transitions in providing a stable re-encoded infectious genome exhibiting *in vivo* attenuation.

#### Impact of “specific” re-encoding strategies

Variants with increased CpG and UpA dinucleotides exhibited attenuated phenotypes *in vivo*, with no mortality, alleviated weight loss and no reduction in viral loads. For variant rUAh (59 UpA), the sequence remained stable throughout the *in vivo* experiments. Hence, the attenuation resulted from the original sequence re-encoding. The originally modified sequence of variant rCGh (59 CpG) remained constant and consensus NS changes were associated with major variants arising during the *in vivo* studies. Such consensus mutations are likely to be adaptive but did not result in complete recovery of virulence during the *in vivo* studies. Altogether, these results are consistent with the attenuating effect of the introduction of CpG/UpA dinucleotides into viral genomes as reported on previous occasions (42, 62-64).

On the other hand, the sequence of the stock variant rTCGh (118 TCG) differed from that of the original construct. To conclude, with respect to the compensatory role of the consensus NS mutations that arose during strain production, further experiments would be required. Nevertheless, the attenuated phenotype (loss of lethality, alleviation of weight loss) of the re-encoded viruses following infection in hamsters suggests a detrimental effect of increasing TCG trinucleotides on YFV virulence *in vivo* that was not completely outweighed by the presence of additional consensus NS changes. Hence, the introduction of TCGs may impact on the *in vivo* phenotype of YFV following infection in vertebrate hosts. Of note, a recent study by Moratorio and colleagues highlighted the adverse effect of the introduction of codons that are likely to yield stop codons (referred to as “1-to-Stop codons”) on the fitness of Influenza A and Coxsackie B3 viruses *in vitro* and *in vivo* (68). Namely, these 1-to-Stop codons are UUA, UUG, UCA and UCG. Only 28 additional UCG were identified within the sequence of YFV rTCG (which was the only re-encoded virus for which there was a change in the number of 1-to-Stop codons). This is 4 to 8-fold lower compared with what was introduced into the genomes of Coxsackie B3 (117 codons) and Influenza A (205 codons) viruses. Nevertheless, further experimentation would be useful to determine whether or not there is an increase in the generation of Stop codons during viral replication of rTCG/rTCGh viruses *in vitro* and/or *in vivo*.

Finally, for strain rNh, consensus NS mutations were identified in the sequences from both the inoculum and hamster liver samples. It is therefore complicated to determine which of the consensus changes were production artefacts or resulted from adaptation of the virus to *in vitro* or *in vivo* conditions. Nevertheless, the virus exhibited a strongly attenuated phenotype *in vivo* (no mortality, alleviated weight loss and viral loads) indicating that, regardless of the emergence of the -potentially compensatory-consensus NS mutations, the combination of several re-encoding strategies led to an important degree of attenuation considering the low number of mutations used for re-encoding (i.e., 233).

#### Adaptation of re-encoded viruses through consensus NS mutations

Altogether, these results indicate that consensus mutations can arise and stably establish in re-encoded viruses in response to *in vivo* conditions. This was observed for most of the re-encoded viruses, notably some with as few as 101 re-encoded sites (rTs1h) or as many as 353 re-encoded sites (rTs3h). In most cases, the consensus mutations did not lead to a full recovery of *in vivo* viral replicative fitness. Observable changes in the biological properties of the viruses (*e.g.* change in tropism, as reported for YF strain 17D), were not reported in any of the viruses that exhibited consensus NS mutations.

#### Immunisation potential of YF-re-encoded strains: serological investigations

For each of the re-encoded viruses that showed complete loss of mortality *in vivo*, sera were recovered from infected hamsters at 16 dpi during the comparative study of pathogenicity (see above). Sera from both uninfected hamsters and vaccinated humans were included as negative and positive controls, respectively. Neutralisation tests were performed using serial fivefold dilutions of all sera, the YF *Ap7M* challenge virus and BHK21 cells. Neutralising antibodies were detected in both hamsters infected with YF-re-encoded viruses (NT_50_ ranging between 2.10^−3^ and 1.10^−5^) and in a control group of 17D-vaccinated humans (NT_50_ between 4 and 5.10^−4^) whilst the negative control group did not show any neutralisation activity against *Ap7M* (see Fig 4).

**Figure 4.**
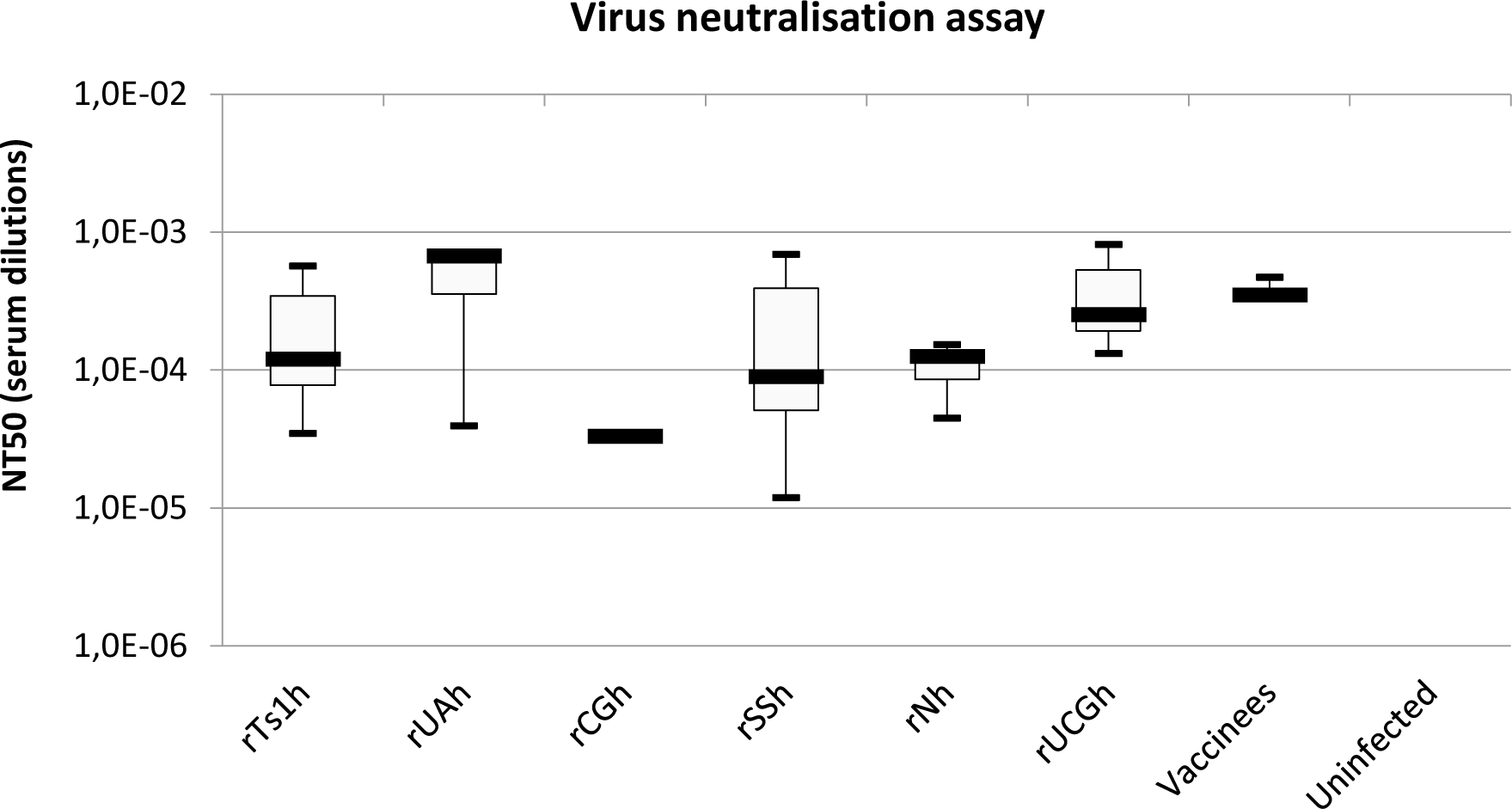
*Results of YFV serology at 16 days post-infection.* Neutralisation assays were performed on sera obtained from hamsters infected with re-encoded viruses that showed complete loss of mortality *in vivo*. For each virus, serum samples were retrieved at 16 dpi from 2 hamsters. Sera from both uninfected hamsters (n=2) and vaccinated humans (vaccinees, n=4) were used as negative and positive controls. Results are expressed as 50% neutralisation titre (NT50) and were evaluated using Graphpad Prism Software (v7.00 for Windows, GraphPad Software, La Jolla California USA, www.graphpad.com). Thick black lines indicate median values and minimum/maximum values are indicated by dashes.

#### Immunisation potential of YF-re-encoded strains: challenge experiments

Two groups of 11 three-week-old female hamsters were inoculated intraperitoneally with 5.10^5^ TCID_50_ of virus (either YFV rCGh or rNh). A negative control group of 9 hamsters was not immunised before the challenge and a control group of 2 uninfected hamsters was included for monitoring hamster weight.

Unexpectedly, one hamster from the YFV rCGh infected group died at 6 dpi, although it did not show any symptom nor weight loss. Its liver was tested positive for YFV. Twenty-four days after inoculation, 2 and 3 hamsters from rCGh and rNh groups, respectively, and a hamster from the non-infected group were euthanised and both serum and liver samples were recovered for neutralisation assay and viral load determination. For all 3 groups, the 8 remaining hamsters were inoculated intraperitoneally with 5.10^5^ TCID_50_ of virus (*Ap7M*). Two hamsters were euthanised at 3 and 6 days post-challenge (dpc) to conduct virology investigations from liver samples whilst 4 were kept for evaluating mortality rate (endpoint in case of survival: 12 dpc). The protection was evaluated by determining for each group (i) viral loads in liver at 3 and 6 dpc, (ii) weight evolution at 6 and 10 dpc, (iii) survival. With regard to the number of samples tested for virology and immunology investigations, no statistical test was achieved in this section. Detailed results for this experiment are available in Fig 5.

**Figure 5.**
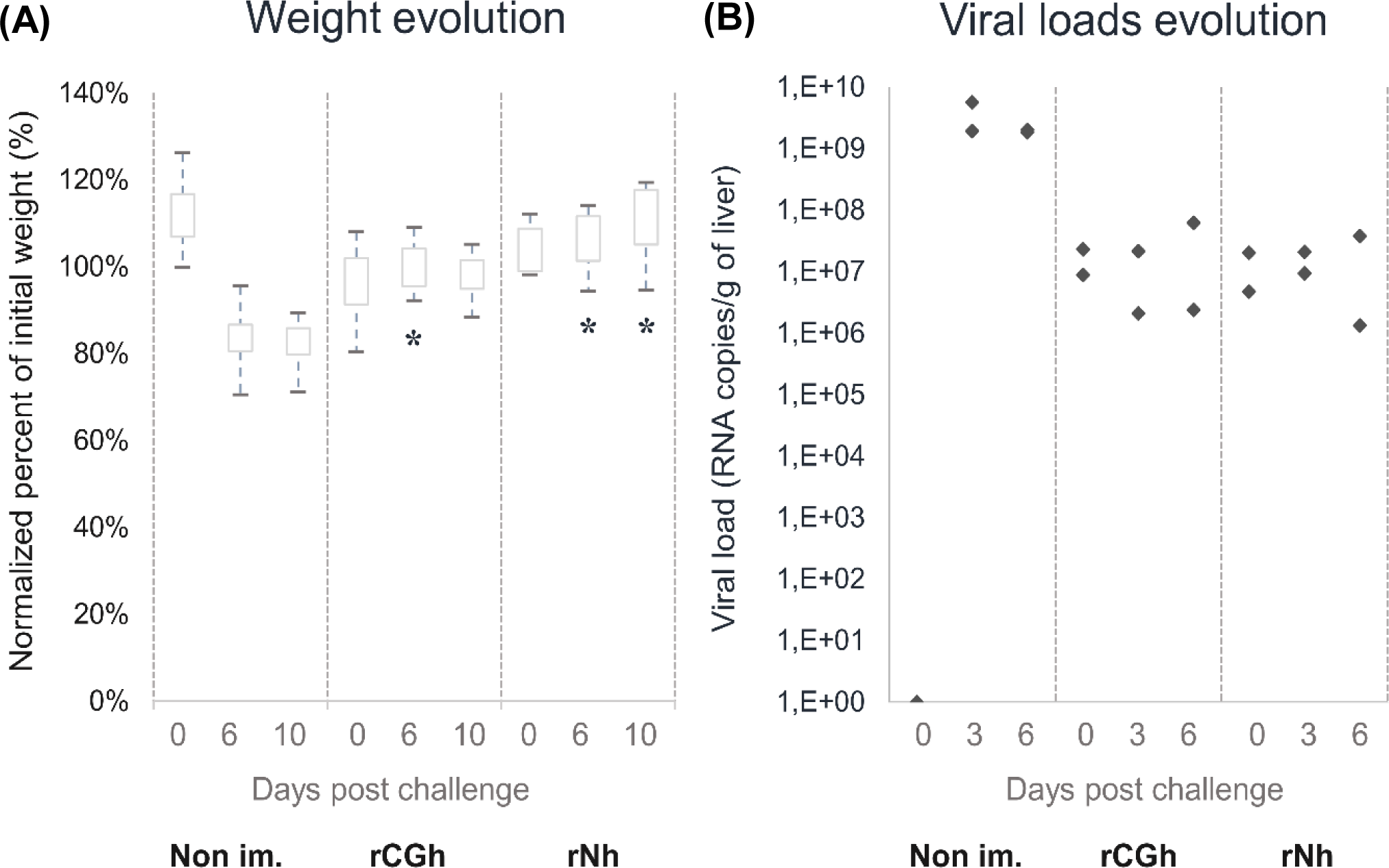
*Challenge assay.* Normalized percent of initial weight (%IW) at day of challenge and 6/10 dpc (A) as well as viral loads at day of challenge and 3/6 dpc (B) are given for all groups (i.e. immunised and control). Viral loads were evaluated on groups of 2 hamsters at 3/6 dpc. %IW were evaluated on groups of 8, 6 and 4 hamsters at 3, 6 and 10 dpc, respectively. Significant %IW differences observed between non-immunised (Non im.) and the other groups at 6/10 dpc (Wilcoxon rank-sum test p-value <0.05) are indicated by a star (*). Viral loads and %IW computations are detailed in the corresponding paragraphs within the Materials and Methods section.

In agreement with previous observations, no mortality was observed post-challenge in either immunised or control groups, indicating that mortality rate in hamsters decreases importantly in adults (>6-week-old) individuals (9). However, there was a great difference in the evolution of viral loads, with no increase in viral yields at 3 and 6 dpc in rCGh and rNh immunised animals while high viral loads were detected in the control animals, around ∼10^9^ RNA copies/g of liver. Evidence of protection was also provided by the fact that there was no weight loss in immunised groups at 6 and 10 dpc, whereas a significant weight loss up to 18% at 10 dpc (Wilcoxon rank sum test p-value=00239) was recorded in the control group. Altogether, these results indicate efficient immunisation of hamsters inoculated with YFV re-encoded strains rCGh or rNh.

## Discussion

Genome re-encoding through synonymous codon replacement is a recognised procedure for attenuating RNA viruses including poliovirus, influenza virus, respiratory syncytial virus, chikungunya virus, tick-borne encephalitis virus and dengue virus (35, 36, 45-47). Additional insights into the mechanisms contributing to attenuation of the virus phenotype following synonymous codon replacement should prove valuable for the custom design of, viable, safe, cost-effective vaccine candidates. The initial purpose of this work was to identify the types of synonymous mutations that would impose the least detrimental effect on viral replicative fitness. Previous codon re-encoding studies in our laboratory were achieved for two other arboviruses, CHIKV and TBEV (45, 46). In the light of this experience, we planned an exploratory study of the effects of re-encoding on both the *in vitro* and *in vivo* phenotypes of yellow fever virus (YFV). Our choice was motivated by the availability of a suitable reverse genetics system for re-encoded strains of YFV production, viz., the ISA method (51)) and a robust YF hamster model (69). Hence, we could test first the *in vitro*, and then the *in vivo* replicative fitness of our strains before using some of the live-attenuated candidates for hamster immunization studies.

Here, we have described the direct impact of synonymous substitutions on YFV replicative phenotype both *in vitro* and *in vivo*. As previously described with TBEV (46), the level of re-encoding necessary to observe viral attenuation *in vivo* was lower than that required to obtain an observable decrease in replicative fitness *in vitro*. As further suggested by supplementary, basic, growth curve experiments (data not shown), all re-encoded strains with only one re-encoded fragment (101-350 mutations) maintained a replicative phenotype close to that of the parental *Asibi* strain *in vitro*. Only YFV rTs4, that included an additional re-encoded fragment (741 mutations in total), showed a severe reduction in replicative fitness *in vitro*. Previous works on Poliovirus, Echovirus, RSV, Influenza, HIV, CHIKV, DENV and Marek’s disease virus (35, 45, 63, 70-74) brought evidence that the effect of synonymous mutations may vary according to the location within the viral sequence. Hence, a control, *Asibi*-derived virus including only the additional re-encoded fragment (388 mutations in the NS5) was produced. Contrarily to the viruses re-encoded in the NS2-3-4 region, it showed a decrease in replicative fitness during competition experiments (outcompeted by the wild-type virus at passage no. 10). As three consensus non-synonymous mutations (associated to major variants) arose in the viral sequence during the production of this virus, it is not possible to associate the *in vitro* phenotype to the re-encoding in the NS5 protein rather than to the additional, consensus, non-synonymous mutations. However, this suggests that in the case of YFV, re-encoding may have a greater effect in the polymerase gene than in other non-structural proteins. In contrast with the *in vitro* results, we observed a broad range of levels of attenuation amongst all the strains that were tested in hamsters (101-350 mutations). Whilst the least attenuated viruses showed no observable loss of replicative fitness and retained a degree of lethality, the most attenuated viruses displayed a complete loss of lethality, together with a tenfold reduction in viral loads in the liver and significant alleviation of weight loss. We did not produce nor test a hamster-adapted version of the rTs4 strain because such a heavily re-encoded virus would have been unlikely to achieve infection *in vivo*.

In contrast with previously published re-encoding studies, we performed deep sequencing of infected hamster liver homogenates to correlate precisely each viral sequence(s) with the observed phenotype. Our results indicate that, detailed genetic information and in particular, that retrieved from *in vivo* experiments, is critical to determine the significance of synonymous substitutions within the attenuation process. When monitoring the evolution of YFV re-encoded strains after production (*i.e.*, transfection followed by 3 passages *in vitro*) or after one passage in hamsters, we observed the spontaneous emergence of both synonymous and non-synonymous substitutions. These consensus changes may be artefacts of the production methods employed (PCR amplification of DNA fragments, stochastic evolution events) or they may have resulted from the adaptation of re-encoded viruses to the differing *in vitro* and *in vivo* conditions of replication. In some cases, we observed the emergence of consensus non-synonymous mutations present in major variants of the viral stock used as the inoculum for *in vivo* experiments. For these viruses, the analysis of the results from *in vivo* experiments and attempts to correlate them with sequencing data was limited due to consensus NS mutations which impeded the identification of the specific impact of silent mutations on the viral phenotype *in vivo*.

Several conclusions can be drawn regarding the *in vivo* phenotypic impact of the different strategies used to produce re-encoded YFV strains.

First, the introduction of simple transitions (*i.e.*, with no modification of putative secondary structures in the genomic RNA) on the replicative fitness of YFV had only minor impact on viral phenotype. They did not cause larger modification of GC%, ENC, Codon Adaptation index (to the homo sapiens codon usage) or in the proportion of codons that are more likely to yield stop codons (“1-to-Stop codons”, as defined by Moratorio and colleagues (68)) than what can be observed amongst clinical/field isolates, indicating that additional mechanisms are probably involved in the attenuation process. Similarly, the introduction of transitions at sites involved in putative genomic secondary RNA structures had a relatively mild detrimental impact on replicative fitness. Second, contrary to secondary RNA structures that have been identified in the untranslated regions of YFV (8) and other flaviviruses (75-77), the putative structures within the virus open reading frame appear unlikely to be important determinants of replicative fitness. Rather, role(s) in finely tuning the efficiency of translation (mRNA or co-translational folding) or in interactions with viral, and non-viral proteins have been proposed (38, 39, 78).

Previous studies have shown that increase in CpG/UpA dinucleotides is an efficient attenuation mechanism (42, 62-64) and this was supported by the evidence of a correlation between increased CpG/UpA and attenuation in our hamster models. However, the introduction of TCG trinucleotides affected the *in vivo* phenotype of YFV. *In silico* analysis predicted that such patterns are likely to be selected against in flaviviruses infecting vertebrates. In addition, studies on influenza virus produced evidence that CpG dinucleotides in an A/U context (e.g., TCG) may enhance virus detection by the host innate immune system (43, 44, 53). Such trinucleotides can be used to induce a host-dependent attenuation, as recently demonstrated for dengue virus (70, 79).

This study approach was exploratory. Hence, we chose to focus on the identification of mutations with low deleterious impact on *in vivo* replication and virulence of the yellow fever virus (YFV) rather than on the comprehensive characterization of the biological mechanisms underlying the attenuation process (e.g., protein translation, RNA replication, activation of innate immunity). Mechanistic insights on the specific effect(s) of the new re-encoding strategies would be most valuable, to gain insight into the finer details of viral attenuation and to facilitate fine-tuning of re-encoded viruses for the specific requirements of their intended use.

The initial aim of this exploratory work was to identify types of mutation with limited detrimental impact on viral replicative fitness, usable for large scale genome re-encoding. In this regard, “non-specific” transitions (*i.e.* with no CpG/UpA creation) represent the mutation type that best met our initial criteria. On this basis, we conclude that the introduction of transitions without CpG/UpA increase may constitute a ground rule for the custom-designed large-scale re-encoding of viral genomes. This does not rule out the possible use of mutations that increase the rate of CpG/UpA dinucleotides. We propose that using a limited number of such mutations may ensure that re-encoded genomes are relatively innocuous when tested *in vivo*, because these patterns ensure enhanced activation of host innate immunity. Thus, procedures combining large, pan-genomic non-specific re-encoding with limited bespoke CpG/UpA re-encoding should lead to the development of safe, stable and effective live-attenuated vaccine viruses with finely tuned phenotypes.

In addition to deciphering the mechanisms of attenuation, evaluating the stability of viral re-encoded variants is a crucial objective for the development of vaccine candidates. Here, we observed no reversion to the parental *Ap7M* sequence of YFV re-encoded viruses when tested *in vivo* and virtually no reversion *in vitro*. However, for several re-encoded strains, the occurrence of consensus non-synonymous mutations in the sequences from the inoculum and/or hamster livers suggests that YFV re-encoded viruses may adapt and evolve both *in vitro* and *in vivo*. The molecular evolution of viral re-encoded strains has already been described for HIV and CHIKV-derived strains *in vitro*. The occurrence of synonymous and non-synonymous consensus mutations correlated with partial recovery of (CHIKV) or full (HIV) WT replicative fitness (45, 72). In accordance with the observations made on CHIKV evolution *in vitro*, some re-encoded strains retained a strongly attenuated phenotype that was not outweighed by the emergence of adaptative consensus mutations. Altogether, these elements suggest that, in some cases, the accumulation of slightly detrimental mutations could trap the virus, with no alternative mutational pathway with which to enable complete restoration of replicative fitness.

As illustrated by the case of poliovirus vaccines, live-attenuated vaccine strains have the potential to evolve following inoculation and in cases of dissemination following vaccination. Hence, the development of re-encoded vaccine candidates should be accompanied by robust studies that address the potential fate of the variants *in vivo*. However, to the best of our knowledge, the evolutionary behaviour of re-encoded viruses *in vivo* has never been studied. Since convenient *in vivo* laboratory models and reverse genetics systems are available, potential live attenuated YFV re-encoded vaccine candidates would provide a convenient experimental model for future *in vivo* studies. In the studies described above, we focused mainly on YF variants re-encoded in one specific region of the genome (NS2A-NS4B). These studies were appropriate for our experimental objectives but would not be ideal for evolution studies. An optimized and more realistic design would rely on *in vivo* serial passage and deep sequencing characterization of evenly distributed synonymous mutations, along the entire genome, each with low fitness impact. The major objective would be to assess the stability of re-encoded strains with synonymously modified codon compositions and in particular, their potential to evolve towards modified phenotypes *in vivo*.

## Methods

*Detailed protocols regarding in silico analysis, virus and cell culture (viral production, transfection and titration (TCID50); competition and virus neutralisation assays), PCR amplification (subgenomic amplicon production), quantitative real-time PCR (qRT-PCR) assays, sample collection and Next-Generation Sequencing (NGS) stand in the supplementary Protocol.*

### Cells and animals

Viruses were produced in Baby hamster kidney BHK21 (ATCC, number CCL10) and Vero (ATCC, CCL81) cells and titrated in BHK21 cells. *In vivo* infection was performed in three-week-old female Syrian Golden hamsters (*Mesocricetus Auratus*, Janvier and Charles River laboratories).

### Ethics statement

Animal protocols were reviewed and approved by the ethics committee “Comité d’éthique en expérimentation animale de Marseille—C2EA—14” (protocol number 2015042015136843-V3 #528). All animal experiments were performed in compliance with French national guidelines and in accordance with the European legislation covering the use of animals for scientific purposes (Directive 210/63/EU).

### Design of re-encoded strains

#### Design of re-encoded strains including only “non-specific” transitions

Re- encoded strains YFV rTs1, rTs2 and rTs3, were designed by introducing increasing numbers of “simple” transitions (siTs) in the CDS of the reference strain *Asibi* (AY640589). The term “simple” refers to synonymous, non-specific (no CpG/UpA introduction) mutations located outside putative secondary structures. Strain YFV rTs4 included all mutations of YFV rTs3 and 388 additional non-specific transitions located between position 6,846 and 9,765 of the CDS, (741 transitions in total).

Strain YFV rSS was re-encoded using 289 siTs as well as 50 non-specific transitions on sites corresponding to putative secondary structures (SII-Ts) identified in the parent sequence (described in the Supplementary Results S2).

### Design of re-encoded strains including “specific” transitions

Viruses YFV rUA and rCG were designed by increasing the number of UpA and CpG dinucleotides in the viral sequence. Transitions leading to UpA dinucleotide introduction (UpA-Ts) were combined with siTs to design strain rUA (59 additional UpA). Similarly, rCG involved a combination of siTs and CpG-Ts (59 additional CpG).

To investigate the impact of introducing TCG patterns on viral replication, mutations allowing the creation of TCG trinucleotides (TCG-Ts) were used for the design of re-encoded strain YFV rTCG (118 additional TCG). Because of the scarcity of editable sites allowing TCG introduction, design rules were relaxed: both transitions and transversions were used and some mutations affected sites corresponding to putative secondary structures (total:12) as well as non-editable sites (total:11).

Finally, a combinatory strategy was used for YFV strain rN design. It included 233 mutations (179 transitions and 54 transversions) that led to the introduction of 83 UpA and 54 CpG dinucleotides into the viral sequence.

### Recovery of infectious viruses and stock production

*All the strains used in this study (1 WT, 1 hamster-adapted and 17 re-encoded) were produced using the ISA method, that enables recovery of infectious viruses after transfection into permissive cells of overlapping subgenomic DNA fragments covering the entire genome (51). All fragment combination schemes used for virus production are detailed in Fig 1 and in the “Fragment combination” section.*

#### Fragments combinations

For *Asibi* strain production, 3 subgenomic fragments were used:FI (positions 1 to 3,919 in CDS and the 5’UTR (119 nucleotides (nt)), FII (3,843 to 6,838) and FIII (6,783 to 10,236 and 3’UTR (509 nt). They correspond to the complete genome of the strain, flanked respectively at 5’ and 3’ termini by the human cytomegalovirus promoter (pCMV) and the hepatitis delta ribozyme followed by the simian virus 40 polyadenylation signal (HDR/SV40pA) (pCMV and HDR/SV40pA sequences were described by Aubry and colleagues (51). The hamster-virulent strain YFV *Ap7M*, was obtained from *Asibi* virus by substituting the FI subgenomic fragment by the FIAp7 fragment, that includes 10 hamster-virulence mutations as described elsewhere (Klitting *et al.*, accepted manuscript).

Re-encoded strains rTs1, rTs2, rTs3, rUA, rCG, rSS, rN and rTCG were derived from *Asibi* by combining a re-encoded version of the 2^nd^ subgenomic fragment FII to the WT 1^st^ (FI) and 3^rd^ (FIII) subgenomic fragments. For the heavily re-encoded rTs4 strain, the WT fragment FI was combined to the re-encoded FII fragment of strain rTs3 and to a re-encoded version of FIII fragment, FIIIR.

All hamster-adapted re-encoded strains (rTs1h, rTs2h, rTs3h, rUAh, rCGh, rSSh, rNh and rTCGh) were obtained by replacing the WT FI fragment with the *Ap7M* FI fragment, FI*Ap7*.

#### Production and recovery

For producing each of the different YFV strains described in this study, 3 overlapping DNA fragments were synthesized *de novo* (Genscript) and amplified by High Fidelity PCR using the Platinum PCR SuperMix High Fidelity kit (Life Technologies) and specific sets of primers. After amplification, all DNA fragments were purified using Monarch PCR & DNA Cleanup kit 5µg (BioLabs) according to the manufacturer’s instructions. Details regarding the subgenomic DNA fragments and the sets of primers used for the amplification step are available in the supplementary Table S6. The different subgenomic fragment combinations are described in Fig 1.

A final amount of 1µg of DNA (equimolar mix of subgenomic cDNA fragments) was transfected using Lipofectamine 3000 (Life Technologies) in a 25 cm2 culture flask of subconfluent cells containing 1mL of culture medium without antibiotics. The cell supernatant media were harvested at 9 days post-transfection (dpi), aliquoted and stored at −80°C. Each virus was then passaged twice in Vero (*Asibi*-derived) or BHK21 (*Ap7M*-derived) cells and once in BHK21 cells. Clarified cell supernatant media from the second passage (virus stocks) were stored and used to perform viral RNA quantification, TCID50 assays and whole-genome sequencing (see corresponding sections).

### Nucleic acid extraction

Samples (liver homogenate or cell culture supernatant medium) were extracted using either EZ1 Biorobot (EZ1 Virus Mini kit v2) or the QiaCube HT device (CadorPathogen kit) both from Qiagen. Inactivation was performed using either 200µL of AVL buffer (EZ1) or 100µL of VXL buffer and 100µL HBSS (Qiacube) according to the manufacturer’s instructions.

### Quantitative real-time RT-PCR assays

All quantitative real-time PCR (qRT-PCR) assays were performed using the EXPRESS SuperScript kit for One-Step qRT-PCR (Invitrogen) and the GoTaq® Probe 1-Step RT-qPCR System (Promega). Primers and probe sequences are detailed in the supplementary Table S6.

The amount of viral RNA was calculated from standard curves using a synthetic RNA transcript (concentrations of 10^7^, 10^6^, 10^5^ and 10^4^ copies/µL). Results were normalised using amplification (qRT-PCR) of the housekeeping gene actin (as described by Piorkowski and colleagues (5)). The standard curves generated for all the YFV-specific assays had coefficients of determination values (R2) >0,98 and amplification efficiencies were between 93% and 100%.

### Competition assays

WT virus was grown in competition with each of the 9, *Asibi*-derived, re-encoded viruses using two RNA ratios in triplicate (WT/re-encoded virus: 10/90 and 50/50). A global estimated MOI of 0.5 was used for the first inoculation. Viruses from each experiment were then passaged nine times with an incubation time of 3 days. At each passage, the estimated MOI was approximately 0.6. After each infection (3 dpi), nucleic acids were extracted from the clarified culture supernatant medium using the QiaCube HT device (see corresponding section) and then used for RT-PCR amplification. The detection of each virus was achieved using next generation sequencing (NGS) methods (see corresponding section), by amplifying a 256 nucleotides region and counting virus-specific patterns within the read population.

### *In vivo* experiments

The laboratory hamster model reproduces the features of human YF but requires the use of adapted strains (7). We recently implemented a hamster model for YFV, based on the use of a strain equivalent to the *Asibi*/hamster p7 strain described by Mac Arthur and colleagues in 2003 (8), *Ap7M*. When inoculated into hamsters, this strain induces a lethal viscerotropic disease similar to that described for YF *Ap7* virus in terms of (i) clinical signs of illness, (ii) weight evolution, (iii) viral loads in the liver and (iv) lethality (100%). This strain was derived from *Asibi* by including 10 mutations in the FI subgenomic region used for viral recovery with the ISA method. As we chose to target the FII subgenomic region for re-encoding, the *Ap7M* strain was well suited for testing the effects of re-encoding on the biological properties of viruses *in vivo*.

Three-week-old female Syrian Golden hamsters were inoculated intra-peritoneally with 5.10^5^ TCID50 of virus in a final volume of 100µL HBSS. In all experiments, a control group of 2 hamsters was kept uninfected. The clinical course of the viral infection was monitored by following (i) the clinical manifestation of the disease (brittle fur, dehydration, prostration and lethargy), (ii) weight evolution and (iii) death. Weight was expressed as a normalised percentage of initial weight (%IW) and calculated as follows:

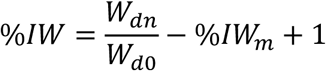

(*W*_*dn*_: weight at day n; *W*_*d0*_: weight on the day of inoculation or challenge; *IW*_*n*_: mean of the *%IW* for control hamsters).

Liver samples were obtained from euthanised hamsters, ground and treated with proteinase K (PK) before nucleic acid extraction using either the EZ1 Biorobot or the QiaCube HT device (see corresponding section). Serum was recovered from euthanised hamsters and stored (- 80°C).

### Virus neutralisation assay

Sera were incubated for 20min at 56°C prior to viral serology. For each serum, serial 5-fold dilutions (first dilution: 1:250) of serum were tested on BHK21 cells for *Ap7M* strain infection inhibition. The plates were incubated for 44 hours before nucleic acid extraction. Virus quantification was achieved using the YFV-specific qRT-PCR system as described above. Then, a viral RNA yield reduction (% of viral inhibition) was calculated for each well as follows:

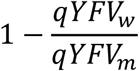

With *qYFV*_*w*_ the number of RNA copies in the analyzed well and qYFV_m_ the mean number of RNA copies in the negative control wells. Fifty percent neutralisation titre NT_50_ values were determined from plots of % viral neutralisation versus serum dilutions and calculated by non-linear regression (four-parameter dose-response curve) analysis using Graphpad PRISM software (v7.00 for Windows, GraphPad Software, La Jolla California USA, www.graphpad.com).

### Whole and partial genome sequencing

Nucleotide sequences were determined using NGS methods: overlapping amplicons spanning either the complete genome sequence or a 256 nucleotides region (nucleotide positions: 4391-4646) were produced from the extracted RNA using the SuperScript® III One-Step RT-PCR System with Platinum®Taq High Fidelity kit (Invitrogen) and specific primers (detailed in the supplementary Table S7). Sequencing was performed using the PGM Ion torrent technology (Thermo Fisher Scientific) following the manufacturer’s instructions.

#### Sequence determination

CDS consensus sequence determination was done using CLC genomics workbench software (CLC bio-Qiagen). Substitutions with a frequency higher than 5% were considered for the analysis of intra-population genetic diversity and major/minor variant subpopulations identification (minor variants: 5%< variants frequency <=75%; major variants: variants frequency >75%). No variant was taken into account with a coverage lower than 1000 reads. No consensus mutation was taken into account with a coverage lower than 50 reads.

#### Re-encoded virus quantification

After sequencing the 256 nucleotide region using the PGM Ion torrent technology. Automated read datasets provided by Torrent software suite 5.0.2 were trimmed according to quality score using CLC genomics workbench software (CLC bio-Qiagen) and 6 re-encoded or wild-type specific patterns (see supplementary Tables S8, S9 and S10) were counted within the read datasets using in-house software. As a control, 2 patterns common to both viruses were counted to evaluate the total amount of virus for each sample.

### Statistical analysis

Viral load comparisons were achieved using Wilcoxon rank-sum test with a continuity correction and Kaplan-Meier survival analysis, using Mandel-Cox’s Logrank tests. Both analyses were performed using R software (6). P-values below 0.05 were considered as significant.

## Aknowledgements

We thank Morgan Seston, Ludivine Molina and Karine Barthélémy from UMR UVE, « Unité des Virus émergents » for their technical assistance. We thank Stéphane Priet from UMR UVE, « Unité des Virus émergents » for his careful re-reading of the manuscript.

## Funding

This work was supported by the European Virus Archive goes Global, http://global.europeanvirus-archive.com/ (European Union’s Horizon 2020 research and innovation programme under grant agreement no. 653316); and the Agence Nationale de la Recherche, http://www.agence-nationalerecherche.fr/ (grant ANR-14CE14-0001 RNA Vacci-Code). The funders had no role in study design, data collection and analysis, decision to publish, or preparation of the manuscript.

## Supporting information

**S1 Supplementary Protocols**

**S2 Supplementary Results**

**S1 Figure. Dinucleotide introduction bias according to the position in the coding frame with-in a 35 YFV CDS alignment.** Mutations were defined with reference to the Asibi strain (Gen-bank accession number: AY640589).

**S2 Figure. Vector/host-preference associated patterns in the *Flavivirus* genus: Comparison of trinucleotides frequencies within the ISF, NKV and DENV groups**

**S1 Table. Yellow Fever dataset: sequence accession numbers**

**S2 Table. Rare codons identified within a complete alignment of 35 YF sequences**

**S3 Table. Putative secondary structures identified within YFV subgenomic fragment FII (position 3924 to 6759)**

**S4 Table. Ecological groups and sequence accession numbers used for vector/host preference associated pattern identification**

**S5 Table. Trinucleotides frequencies amongst ISF, MBF and NKV groups.** Each frequency was expressed as the ratio between the trinucleotide count and the total number of trinucleotides in the considered sequence. The differences in frequencies were then calculated and summed between ISF and both MBF and NKV groups and between MBF and both NKV and ISF groups.

**S6 Table. PCR/qRT-PCR primers and Yellow fever subgenomic fragment description**

**S7 Table. RT-PCR primers used for pre-sequencing amplification**

**S8 Table. Universal, wild-type and re-encoded-specific patterns (1)**

**S9 Table. Universal, wild-type and re-encoded-specific patterns (2)**

**S10 Table. Universal, wild-type and re-encoded-specific patterns (3)**

**S11 Table. Mature proteins positions within Yellow Fever Virus polyprotein sequence.** (obtained from UniProt database, www.uniprot.org)

